# Serotonin Modulates an Inhibitory Input to the Central Amygdala from the Ventral Periaqueductal Gray

**DOI:** 10.1101/2022.03.28.486055

**Authors:** Olivia J. Hon, Jeffrey F. DiBerto, Christopher M. Mazzone, Jonathan Sugam, Daniel W. Bloodgood, J. Andrew Hardaway, Mariya Husain, Alexis Kendra, Nora M. McCall, Alberto J. Lopez, Thomas L. Kash, Emily G. Lowery-Gionta

**Affiliations:** Bowles Center for Alcohol Studies, School of Medicine, University of North Carolina at Chapel Hill; Department of Pharmacology, School of Medicine, University of North Carolina at Chapel Hill

## Abstract

Fear is an adaptive state that drives defensive behavioral responses to specific and imminent threats. The central nucleus of the amygdala (CeA) is a critical site of adaptations that are required for the acquisition and expression of fear, in part due to alterations in the activity of inputs to the CeA. Here, we characterize a novel GABAergic input to the CeA from the ventral periaqueductal gray area (vPAG) using fiber photometry and *ex vivo* whole-cell slice electrophysiology combined with optogenetics and pharmacology. GABA transmission from this ascending vPAG-CeA input was enhanced by bath application of serotonin via activation of serotonin type 2C (5HT_2C_) receptors. Results indicate that these receptors are presynaptic. Interestingly, we found that GABA release from the vPAG-CeA input is enhanced following fear learning via activation of 5HT_2C_ receptors and that this pathway is dynamically engaged during fear learning. Additionally, we characterized serotonin release in the CeA during fear learning and recall for the first time using fiber photometry coupled to a serotonin biosensor. Together, these findings describe a mechanism by which serotonin modulates GABA release from ascending vPAG GABA inputs to the CeA and characterize a role for this pathway in fear learning.

## INTRODUCTION

In canonical fear models, sensory information from thalamic and cortical areas is conveyed to the lateral and basolateral amygdala [1–4]. Adaptations within these subregions encode associations between fear-inducing stimuli and environmental cues, and are thought to form the neurobiological basis for fear learning. Information from the lateral and basolateral amygdala is then conveyed to the central amygdala (CeA), the major output nucleus of the amygdala. The CeA governs conditioned fear responses through projections to output structures, namely the periaqueductal gray (PAG)[5].

Disruption of CeA activity impairs fear learning and expression, while stimulation of the PAG elicits defensive responses and disruptions of PAG activity impair conditioned freezing responses [6–10]. Recently, evidence of plasticity in the CeA following fear learning has broadened the view that adaptations throughout the amygdala underlie fear learning. This plasticity has been observed within the complex inhibitory microcircuits of the CeA, as well as at CeA inputs arising from ‘upstream’ regions of the fear circuit like the thalamus [9,11–14]. However, mounting evidence indicates that CeA inputs arising from ‘downstream’ effector targets may also influence fear encoding. Recent studies support a role for the PAG in the integration and processing of fear-eliciting stimuli [10,15–17], and that the PAG modulates ‘upstream’ amygdalar subregions like the basolateral and lateral amygdala [10,15]. Thus, outputs from the PAG to the amygdala may play a critical role in fear encoding.

The CeA receives inputs from the ventral PAG (vPAG) [18,19], though the molecular phenotype of this input has not been characterized. The vPAG contains diverse neuronal populations, among the largest of which are GABA neurons [20,21], and converging lines of evidence suggest a role for vPAG GABA neurons in fear learning. Recent work from our lab shows that chemogenetic inhibition of vPAG GABA neurons during fear learning disrupts subsequent fear expression [22]. Others have shown that optogenetic activation and inhibition of vPAG GABA neurons bidirectionally modulate freezing behavior [23], highlighting a key role for this population in fear learning.

Serotonin (5-hydroxy tryptamine, 5HT) has been implicated in the modulation of fear learning [24–26], and selective serotonin reuptake inhibitors are the first-line treatment for anxiety and mood disorders. However, the precise mechanisms through which 5HT exerts its effects on fear are unknown. We previously showed that 5HT is released in both cortical and amygdalar regions during fear learning [27], while others have shown that 5HT neurons in the dorsal raphe nucleus that project to the CeA are responsive to footshock stimuli [28]. Importantly, 5HT modulates GABAergic transmission in many hubs for affective behaviors [29–31], including the CeA [32], raising the possibility that 5HT modulation of GABA in the CeA is important for fear encoding.

Here, we demonstrate that vPAG GABA neurons directly project to the CeA to modulate activity and influence fear learning using fiber photometry and *ex vivo* slice electrophysiology paired with optogenetics and pharmacology. We characterize 5HT release in the CeA and assess 5HT modulation of this pathway and its role in fear-induced plasticity.

## METHODS

### ANIMALS

*Vgat*-ires-Cre [33] mice obtained from Dr. Bradford Lowell were used for all input-specific experiments to allow specific targeting of GABA neurons. *Vgat-*ires-Cre mice were bred in house from *Vgat*-ires-Cre x C57BL6/J (Jackson Laboratories, Bar Harbor, ME) pairings. Male *Vgat*-ires-Cre mice >8 weeks of age were used for all experiments. To characterize 5HT’s effects on GABA transmission in the greater CeA population and 5HT release in the CeA, male C57BL6/J mice were used. All mice were maintained on a 12h/12h light/dark cycle with lights on at 7 AM. Mice had access to standard rodent chow and water *ad libitum* for the duration of experiments. All experiments were conducted in accordance with the University of North Carolina at Chapel Hill’s Institutional Animal Care and Use Committee’s guidelines.

### SURGERY

Mice were deeply anesthetized using isoflurane (4% for induction, 1-2% for maintenance). Mice were administered buprenorphine (.1 mg/kg, s.c.) during surgery, and were also provided with an acetaminophen solution in the home cage (80 mg/200mL in water) one day before and for at least 3 days following surgery. AAV constructs were delivered to the vPAG (AP −4.6mm, ML 0.0, DV-3.2mm, 20° angle, 0.5 µl) or CeA (AP −1 mm, ML 2.8, DV −4.7, 0.3 µl) at a rate of 0.1 µl/ min using Hamilton Neuros microsyringes. For photometry experiments, optical fibers were implanted at the same coordinates as virus delivery and cemented to the skull with dental cement (C&B Metabond, Parkell). Mice recovered for at least 6 weeks prior to the start of experiments to allow for expression of the viral constructs at terminals.

### FEAR CONDITIONING

Cued fear conditioning was performed using a three-day protocol. On the first day (fear learning), mice were placed into a fear conditioning chamber with a shock grid floor (context A, Med Associates) cleaned with a 20% ethanol + 1% vanilla extract solution. Following a 2 minute baseline, 5 tone-shock pairings (tone: 30 s, 80 dB, 3 KHz, shock: 0.6 mA, 2 s) were presented, separated by a random inter-trial interval of 60-90 seconds. Naïve mice received tones but not shocks. On the second day (context recall), mice were tethered for fiber photometry recordings and placed into an empty cage for 5 minutes. Mice were then placed back into context A for 15 minutes. On the third day, mice were placed into a novel context (context B) with white plastic flooring, curved white walls, and cleaned with a 0.5% acetic acid solution. Following a 2 minute baseline, 5 tones (identical to fear learning) were presented separated by a random ITI of 60-90 s. Behavior hardware and video recording was controlled by Ethovision XT software (Noldus Inc.). Mice for electrophysiology experiments only underwent fear learning, mice for iSeroSnFR recordings underwent fear learning and then cued recall 24h later, and mice for GCaMP recordings underwent the full 3-day protocol.

### BEHAVIOR TRACKING AND ANALYSIS

Behavior was tracked and scored for freezing using open-source, machine learning platforms. First, videos were tracked using DeepLabCut software that uses deep neural networks to estimate mouse position [34]. Tracking information was then analyzed using SimBA, a machine learning platform that can be trained to identify and quantify specific behaviors like freezing [35]. A freezing classifier was developed in-house and then refined such that it did not differ by more than 10% from hand-scoring. Freezing data were then processed using a custom MATLAB (Mathworks) script to calculate the % freezing during different epochs of behavior trials, and the start and end time of each freezing bout.

### FIBER PHOTOMETRY

For all experiments, optical fibers (200 µm diameter, 0.37 NA, Newdoon Technologies, Hangzhou, China) were tested for light transmission prior to implantation. Prior to the start of experiments, mice were habituated daily to patch cord tethering for 3 days. Data were recorded using a Neurophotometrics FP3001 system (Neurophotometrics, San Diego, CA). Briefly, 470 nm and 415 nm LED light was bandpass filtered, reflected by a dichroic mirror, and focused onto a multi-branch patch cord (Doric, Quebec City, Quebec) by a 20x objective lens. For all recordings, alternating pulses of 470 nm and 415 nm light were delivered at ∼50 µW at a rate of 40 Hz. Photometry signals were time-locked to behavior using a custom built Arduino designed to digitize TTL timestamps from behavior hardware. Signals were analyzed using custom MATLAB scripts as detailed below.

#### GCaMP in vPAG^VGAT^ neurons

GCaMP7f (AAV8-syn-FLEX-jGCaMP7f-WPRE, Addgene) was injected into the vPAG of *Vgat*-ires-Cre mice and fibers were implanted in vPAG and bilaterally in the CeA. 470 and 415 signals were deinterleaved, and background fluorescence was subtracted. Signals were then lowpass filtered at 5 Hz and fit to a biexponential curve, and the fit was subtracted from each trace to correct for baseline drift. dF/F for both 470 and 415 signals was calculated as (raw signal-fitted signal)/(fitted signal), and dF/F traces were then z-scored. The 415 signal was then fit to 470 signal using non-negative robust linear regression to correct for motion [36]. For spike analysis: First, the mean absolute deviation (MAD) of the baseline period (first minute of each recording) was calculated. vPAG GCaMP spikes were then identified using a custom MATLAB script. Peaks with a prominence greater than 2*MAD were included in analysis. Alignment with freezing behavior: Freezing behavior was first processed as described above. For vPAG spiking: the timing of each peak was then aligned with frame-by-frame freezing data to determine whether each spike occurred during freezing or mobility. For CeA: the start and end time of each freezing bout was extracted. CeA data was segmented based on these timestamps and the average z-score was calculated across all freezing and mobility bouts.

#### iSeroSnFR in CeA

iSeroSnFR (AAV9-CAG-iSeroSnFR-nlgn, plasmids provided by Dr. Lin Tian, AAV produced at UNC Vector Core, Chapel Hill, NC) or AAV9-hSyn-EGFP (diluted 1:10 in sterile PBS, Addgene) was injected bilaterally into the CeA of C57Bl/6J mice. 470 and 415 signals were deinterleaved, and background fluorescence was subtracted. 470 signals were lowpass filtered at 5 Hz and fit to a bioexponential curve, and the fit was subtracted from each trace to correct for baseline drift. dF/F was calculated as (raw signal-fitted signal)/(fitted signal), and dF/F traces were then z-scored. 415 signal was not used in iSeroSnFR analysis as it not an appropriate wavelength for motion control for this sensor. Peri-event plots were generated by normalizing traces to the first 20 seconds of each trace.

### HISTOLOGY

For photometry experiments, mice were transcardially perfused 30mls each of phosphate buffered saline and 4% paraformaldehyde. Optical fibers were left in the brain for 3 hours prior to brain extraction. Brains were post-fixed in 4% PFA for 24h, then sectioned at 45 µm on a vibratome. GCaMP and iSeroSnFR fluorescence was amplified using immunohistochemistry. Briefly, free-floating sections were washed 3x 5min in PBS, once in 0.5% Triton-X in PBS, and then 3x 5min more in PBS. Slices were then incubated in a blocking solution made with 10% normal donkey serum, 0.1% triton-X in PBS for 1 hour. Slices were then transferred into a primary antibody solution made with chicken anti-GFP IgY (Aves Labs, Tigard, OR) diluted 1:500 in blocking solution, and incubated overnight. The following day, sections were washed 3x 10 min in PBS, and then incubated in a secondary antibody solution made with donkey anti-chicken IgY conjugated to Alexa-Fluor 488 (Jackson Immunoresearch, West Grove, PA) diluted 1:200 in PBS for 2 hours. Finally, sections were washed 4x 10 min in PBS. Sections were imaged on a fluorescence microscope, and viral expression and fiber placement were verified. Mice with off-target expression or placement were excluded from analysis.

### ELECTROPHYSIOLOGY

Whole cell patch clamp recordings were performed in normal artificial cerebral spinal fluid (ACSF; in mmol/L: 124 NaCl, 4.4 KCl, 2 CaCl_2_, 1.2 MgSO_4_, 1 NaPO_4_, 10 glucose, 26 NaHCO_3_) on brain slices prepared from mice sacrificed via deep isoflurane anesthesia as previously described [37,38].

#### vPAG^VGAT^-CeA Specific Transmission

*Vgat*-ires-Cre mice were injected with ChR2 in the vPAG and slices containing the vPAG were prepared. Whole-cell patch clamp slice electrophysiology was conducted to characterize the firing of action potentials in response to 1-ms pulses of 490 nM blue light in vPAG GABA neurons at increasing frequencies (1-100 Hz) using a potassium-gluconate based intracellular solution (in mM: 135 K+ gluconate, 5 NaCl, 2 MgCl2, 10 HEPES, 0.6 EGTA, 4 Na2ATP, 0.4 Na2GTP, pH 7.3, 285-290 mOsmol). In slices containing the CeA, slice electrophysiology was conducted to characterize inhibitory post-synaptic currents (IPSCs) evoked by 1-10 ms pulses of 490 nM blue light (Cool LED, Traverse City, MI) in CeA cells proximal to vPAG GABA inputs. In the CeA, light-evoked IPSCs (eIPSCs) were isolated by adding the glutamate receptor antagonist kynurenic acid (3 mM; Kyn) to ACSF and using a potassium-chloride/potassium-gluconate based intracellular solution (in mmol/L: 70 KCl, 65 K+-gluconate, 5 NaCl, 10 HEPES, 0.6 EGTA, 4 adenosine triphosphate, 0.4 guanosine triphosphate, pH 7.2, 290 mOsmol). Bath application of the GABA-A receptor channel blocker picrotoxin (25 µM) was used to confirm that evoked events were GABAergic. A paired-pulse protocol consisting of 2 × 1-10 ms pulses of 490 nM blue light separated by a 50 ms inter-pulse interval was used to further investigate the pathway. To evaluate the ratio between the amplitude of the second pulse relative to the amplitude of the first pulse (termed the paired pulse ratio or PPR), the intensity of the LED-driven light pulse was adjusted for each cell to reliably elicit 2 time-locked eIPSCs.

#### Serotonin Effects on vPAG^VGAT^-CeA-Specific Transmission

Following establishment of a stable baseline, 5HT (10 µM) was bath applied for 10 minutes, followed by a 10 minute washout period, at a flow rate of 2 ml/min. This procedure was also used with the sodium channel blocker tetrodotoxin (TTX; 1 µM) and the voltage-dependent potassium-channel blocker 4-Aminopyridine (4-AP; 100 µM) in the bath in addition to kynurenic acid. To determine if 5HT_2C_ receptors mediate the effect of serotonin on this input, 5HT_2C_ receptor antagonist RS102221 (5 µM, Tocris, Minneapolis, MN) was added in the bath. For all experiments, slices pre-bathed in ACSF + drug solutions for at least 20 minutes prior to the start of recording.

#### Serotonin Effects on Spontaneous and Miniature GABA Transmission in a Broad Medial CeA Population

Whole-cell recordings from the CeA were conducted to assess the effects of 5HT on spontaneous network-dependent inhibitory post-synaptic currents (sIPSCs) using kynurenic acid (3mM) in the bath solution and a potassium-chloride/potassium-gluconate based intracellular solution to isolate GABAergic transmission. After a 2 minute baseline, 5HT (10 µM) was bath applied as described above. The effects of 5HT on miniature network-independent inhibitory post-synaptic currents (mIPSCs) were assessed using identical parameters as those for sIPSCs with the addition of TTX (1 µM) to the bath solution. The role of 5HT_2C_ receptors in 5HT effects on sIPSCs was assessed as described for sIPSC experiments except that slices were pre-bathed in the 5HT_2C_ receptor antagonist RS102221 (5 µM). Presynaptic and postsynaptic effectss of 5HT on GABA release were assessed by analyzing IPSCs frequency and amplitude, respectively. Responses to 5HT were defined as a change of greater than 50% of baseline values. Any response that was within 50% of baseline values was classified as no change.

#### vPAG^VGAT^-CeA Specific Transmission following Fear Learning

To test if the vPAG-CeA pathway undergoes plasticity during fear learning, mice underwent fear learning as described above. 15 minutes after the end of the session, mice were sacrificed. Using the PPR protocol, basal GABA release from this pathway, as well as serotonin’s ability to modulate release from this pathway were evaluated. In addition, the ability of 5HT_2C_ receptors to modulate GABA release from this pathway following fear learning was investigated by administering the 5HT_2C_ receptor antagonist SB242084 (Tocris, Minneapolis, MN. 3 mg/kg in 10% cremophor + 0.9% saline in a 10 ml/kg i.p.) 30 minutes prior to the fear learning session (shock group) or 45 minutes prior to sacrifice (naïve group).

### STATISTICS

Sample sizes for all experiments are based on those commonly used in the literature to assess modulatory effects or between-group differences in slice electrophysiology and behavior experiments. Group numbers are provided in the figure captions. Paired two-tailed t-tests were used to assess the effects of 5HT by comparing the final four minutes of 5HT wash on (minutes 11-14) or the final four minutes of washout (minutes 20-23) to the baseline values (minutes 0-4). Between group differences were assessed using unpaired t-tests, two-way ANOVA (treatment x drug) or repeated measures ANOVA with treatment (naïve vs. shock) and time as factors. Where indicated, *a priori* hypotheses were tested using planned comparison unpaired two-tailed t-tests. Statistical significance was accepted at p-values <0.05.

## RESULTS

### The CeA receives GABAergic inputs from the ventral PAG (vPAG)

To determine if there is a GABAergic projection from vPAG to CeA, we used whole-cell slice electrophysiology combined with optogenetic stimulation of Cre-inducible ChR2 injected in the vPAG of *Vgat*-ires-Cre mice. We first confirmed that vPAG^VGAT^ neurons expressing ChR2 are activated in response to 1-ms pulses of 490 nM light. We found that vPAG^VGAT^ neurons fire action potentials with 100% fidelity in response to light pulses up to 10 Hz (Fig. 1 E-G). To explore the possibility that the CeA receives GABA inputs from the vPAG, we used ChR2-assisted circuit mapping. We discovered terminals in the CeA originating from vPAG GABA neurons, uncovering a dense nexus of synapses formed between vPAG GABA neurons and neurons of the CeA, particularly in the medial CeA (see Fig. 1C-D). Notably, terminals were not visualized in the adjacent basolateral amygdala (BLA) which receives inputs from the dorsal PAG [10]. We functionally confirmed our anatomical observation by light-evoking eIPSCs in the CeA. eIPSCs were GABA-A receptor mediated, as they were blocked in the presence of picrotoxin but not in the presence of kynurenic acid (Fig. 1H). eIPSCs were detected in 56.86% of cells recorded from the CeA (Fig. 1I). Importantly, this response was time-locked to the light pulse and persisted in the presence of TTX and 4-AP, indicative of a monosynaptic response.

**Figure 1.**
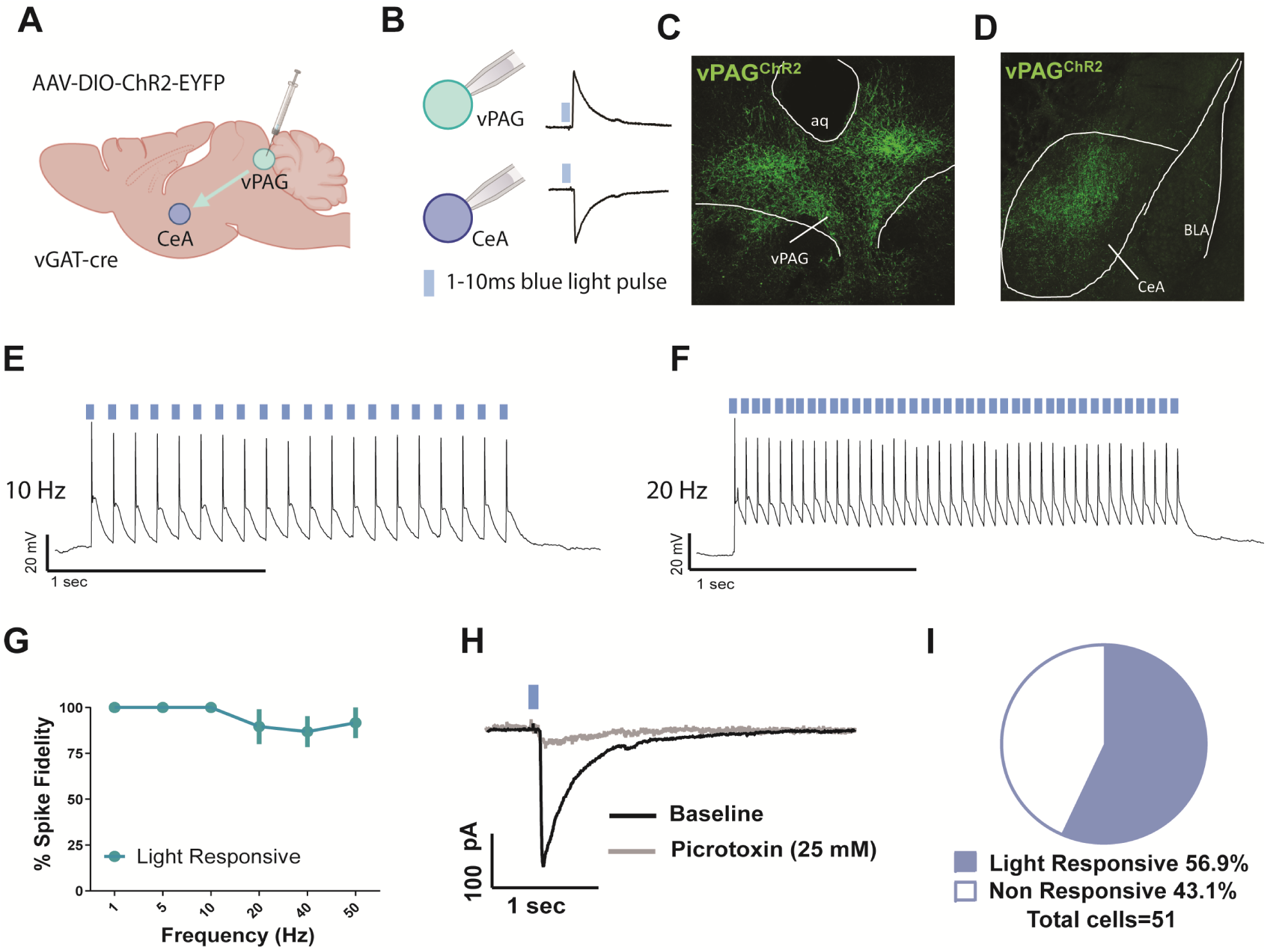
vPAG GABA neurons have functional projections to the CeA. **A:** surgical schematic of channelrhodopsin infusion into the vPAG of *Vgat*-ires-Cre mice. **B:** schematic of responses in vPAG and CeA upon blue light stimulation. **C, D:** Representative image showing expression of channelrhodopsin (ChR2) in the vPAG of a *Vgat*-ires-Cre mouse. **E, F:** Representative action potential firing at 10 Hz and 20 Hz. **G**: Fidelity of action potential spiking in response to ChR2 activation by 1 ms pulses of blue 490 nM light in vPAG GABA neurons (c; n= 6 cells from 5 mice). **H:** representative average trace of inhibitory post-synaptic potentials evoked by 1 ms pulses of blue 490 nM light (eIPSCs) recorded from a cell of the CeA proximal to ChR2 terminal expression in the absence and presence of the GABA-A ion channel blocker picrotoxin (25 nM). **I:** Percent of cells recorded from in the CeA proximal to ChR2 terminal expression that showed eIPSCs in response to blue 490 nM light stimulation (n= 51 cells).

### Serotonin is released in the CeA during fear learning

5HT neurons in the dorsal raphe that project to the CeA are responsive to footshock [28], and we have shown previously that 5HT is released into other subregions of the amygdala during fear learning [27]. Thus, we hypothesized that 5HT is also released in the CeA during fear learning. Using fiber photometry coupled to a genetically encoded 5HT sensor, iSeroSnFR [27], we recorded 5HT dynamics bilaterally in the CeA (Fig. 2A,B). We found that extracellular 5HT decreases sharply during shock delivery, then rebounds above baseline, peaking roughly 20 seconds after shock. Both shock and post-shock responses were significantly different from naïve (iSeroSnFR, no shock), and GFP (GFP, shock) controls [shock response: F=10.94 p=.0003 one way ANOVA, iSeroSnFR fear v. naïve p=.0014, iSeroSnFR fear v. GFP p=.0019, post-shock: F=12.32 p=.0001, iSeroSnFR fear v. naïve p=.0004, iSeroSnFR fear v. GFP p=.0023] (Fig 2G-K). Notably, 5HT dynamics did not differ between hemispheres (data not shown). Additionally, iSeroSnFR fear mice froze significantly more than naïve mice, indicating that the 5HT sensor did not impair fear learning [F(1,22)=4.846 p=.0385 two way ANOVA] (Fig. 2F). Comparison of responses during the first and last tone-shock pairing revealed changes in 5HT release over the course of fear learning. In iSeroSnFR fear mice, the post-shock response significantly decreased between the first and last presentations [t(12)=2.431 p=.0317 paired t-test] (Fig 2L-M). No changes were detected in iSeroSnFR naïve or GFP shock mice, indicating that this change is likely physiological and not due to photobleaching. One day following fear learning, fear expression was tested in a cued recall test and 5HT dynamics were recorded. Responses to tone presentations did not differ statistically between groups (Fig. S1C-G).

**Figure 2.**
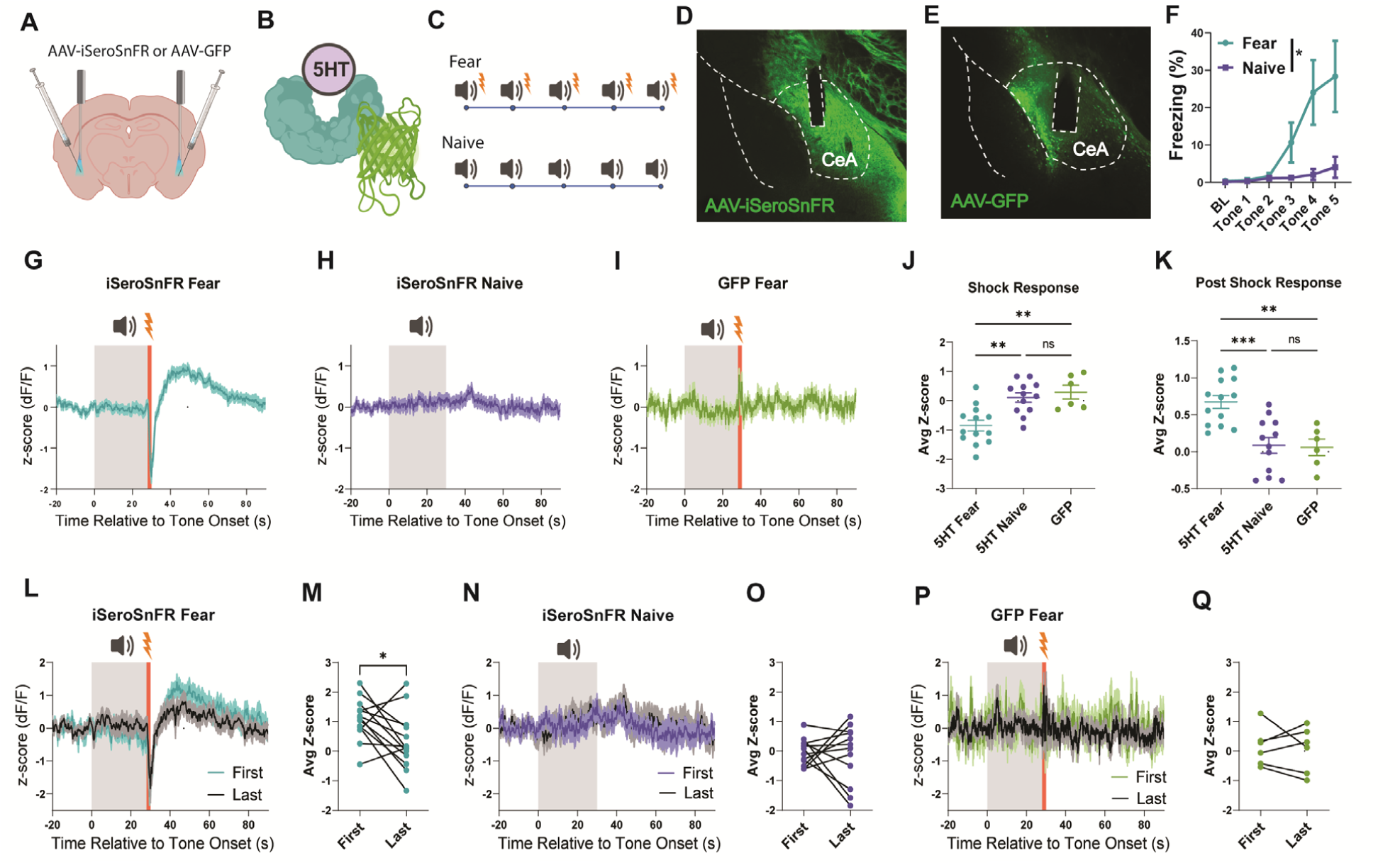
5HT is released in the CeA during fear conditioning. **A:** schematic of injections and fiber implant for iSeroSnFR recordings. **B:** schematic of iSeroSnFR. **C:** Schematic of tones and shocks presented during fear learning for fear and naïve groups. **D, E:** Representative image of iSeroSnFR (D) and GFP (E) and optic fiber placement in CeA. **F:** Freezing behavior during fear learning. **G-I:** response to tone and shock presentations averaged across cohort for iSeroSnFR Fear (G), iSeroSnFR Naïve (H), and GFP Fear (I). **J:** shock response averaged from t=28-30 relative to tone onset across all trials. **K:** post-shock response averaged from t=33-60 relative to tone onset across all trials. **L, N, P:** responses to first and last tone/shock presentations averaged across cohort for iSeroSnFR Fear (L), iSeroSnFR Naive (N), and GFP fear (P). **M, O, Q:** average post-shock response for first and last trials for iSeroSnFR Fear (M), iSeroSnFR Naive (O), and GFP fear (Q). n= 13 iSeroSnFR fear, 11 iSeroSnFR naïve, 6 GFP fear

### The vPAG^VGAT^-CeA input is modulated by 5HT directly and via presynaptic 5HT_2C_ receptors

5HT has been shown previously to modulate GABAergic transmission in the CeA [32], thus we postulated that 5HT may also control GABA release from vPAG^VGAT^ terminals. To explore the effect of 5HT on GABA release specifically from the vPAG^VGAT^-CeA input, we expressed Cre-inducible ChR2 in the vPAG of *Vgat*-ires-Cre mice and recorded eIPSCs evoked by a 1 × 1-10 ms pulse of light or eIPSCs evoked by 2 × 1-10 ms pulses of light separated by a 50 ms inter-pulse interval in the CeA. In all cells tested, 5HT significantly enhanced the amplitude of the first eIPSC [t(12)= 4.562, p= 0.0007 for baseline v. 5HT; Fig. 3 B-D], an effect that did not wash out [t(12)= 3.539, p= 0.0041 for baseline v. washout; data not shown], demonstrating that 5HT enhances GABA transmission at vPAG synapses with CeA neurons. In addition, 5HT significantly reduced the paired-pulse ratio (PPR) [t(5)= 2.817, p= 0.0372 for baseline v. 5HT; t(5)= 4.202, p= 0.0085 for baseline v. washout; see Fig. 3E], indicating that 5HT enhances GABA release at vPAG synapses on to CeA neurons presynaptically. To determine if this effect is direct, we isolated direct transmission by adding TTX and 4-AP to the bathing solution. We found that the ability of 5HT to enhance GABA release persisted in the presence of TTX and 4-AP [t(3)= 10.98, p= 0.0016 for amplitude of the first peak baseline v. 5HT; t(3)= 6.577, p= 0.0072 for PPR baseline v. 5HT], demonstrating that 5HT acts directly at the vPAG^VGAT^-CeA synapses to enhance GABA release (Fig. 3F-I). The 5HT_2C_ receptor subtype is Gq-coupled and modulates GABA release in the amygdala [31,32,39], and so we surmised that it may mediate 5HT effects on GABA release from vPAG terminals. We found that the ability of 5HT to enhance GABA release from the vPAG^VGAT^-CeA input was blocked in the presence of the 5HT_2C_ antagonist RS102221 [t(4)= 1.122, p= 0.3248 for amplitude of the first peak baseline v. 5HT; t(4)= 1.477, p= 0.2137 for PPR baseline v. 5HT; see Fig. 3 J-M], supporting the idea that presynaptic 5HT_2C_ receptors potentiate GABA release in this pathway.

**Figure 3.**
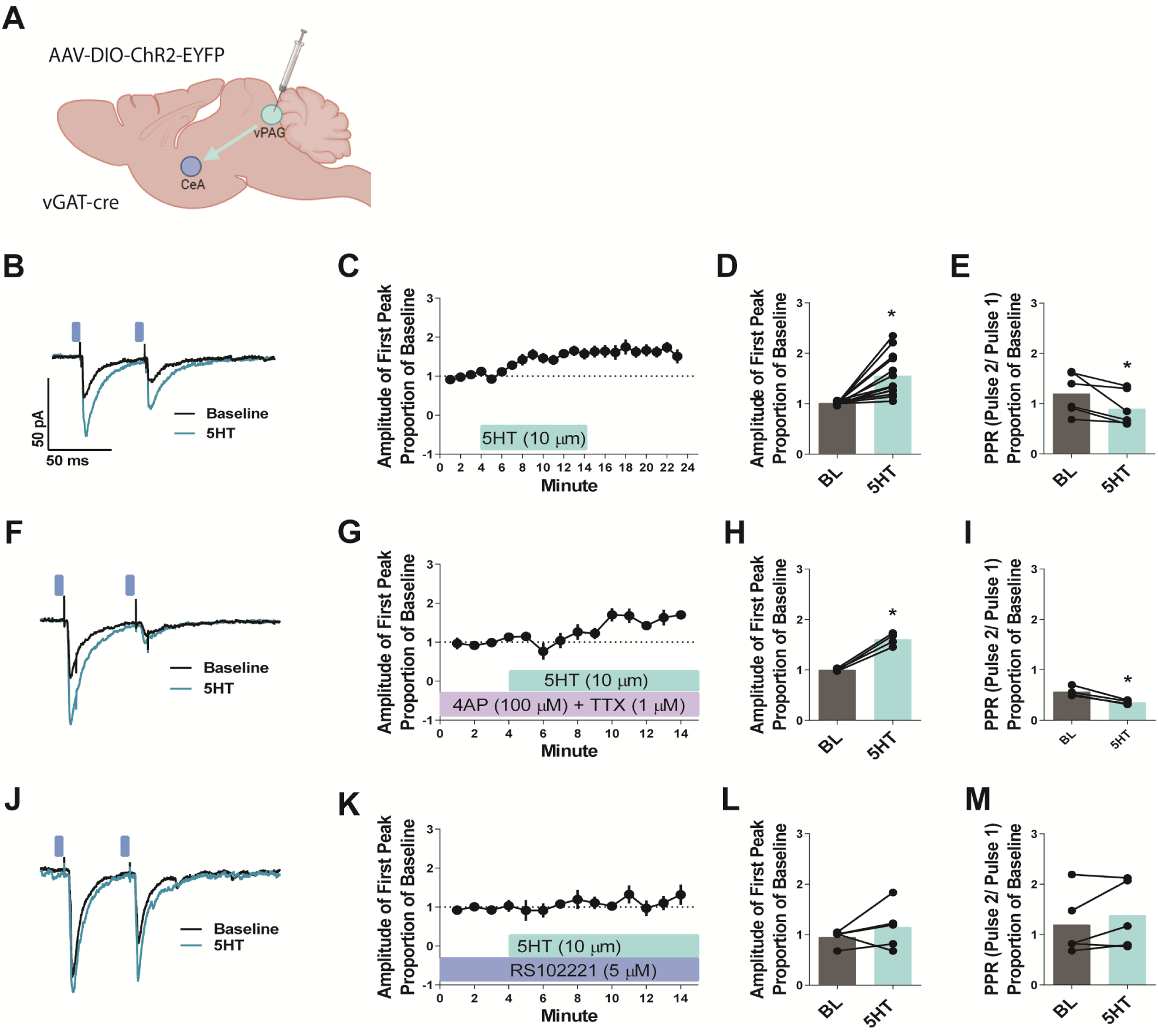
The vPAG^Vgat^-CeA pathway is modulated by serotonin directly and via presynaptic 5HT2C receptors. **A:** surgical schematic of channelrhodopsin infusion into the vPAG of *Vgat*-ires-Cre mice. **B:** Representative traces showing eIPSCs at baseline and during serotonin bath application. **C-D:** Time course and summary of the effects of bath application of serotonin (10 µM) on the amplitude of the first evoked peak relative to baseline values (n= 13 cells from 9 mice). **E**: Summary of the effects of serotonin on the paired pulse ratio (PPR; amplitude of pulse 2/ amplitude of pulse 1) (n= 6 cells from 4 mice). **F-I:** Time course and summary of the effects of serotonin on the amplitude of the first evoked peak (F, G) and PPR (H, I) in the presence of 4-Aminopyridine (4 AP; 100 µM) and tetrodotoxin (TTX; 1 µM) (n= 4 cells from 2 mice). **J-M:** Time course and summary of the effects of serotonin on the amplitude of the first evoked peak (J,K) and PPR (L,M) in the presence of the 5HT2C antagonist RS102221 (5 µM) (n= 5 cells from 4 mice). * denotes *p*< 0.05; all data shown as means +/- SEM.

To explore whether these effects were consistent across the general medial CeA, we recorded spontaneous IPSCs in the CeA of C57BL/6J mice. 5HT (10 µM) enhanced the frequency [p<.0465 t(10)=2.271] (Fig. S2 B-C), but not the amplitude (Fig. S2 A, C) of sIPSCs in 63.6% of recorded cells (Fig. S2 D) suggesting that overall 5HT enhances presynaptic GABA release in the medial CeA. 5HT enhanced the frequency but not amplitude of mini IPSCs (mIPSCs) recorded in the presence of TTX (1 µM) only in a subset of cells (Fig. S2 E-H), suggesting that the large majority of this effect appears to be polysynaptic. 5HT_2C_ antagonist RS102221 partially blocked 5HT enhancement of sIPSC frequency in the medial CeA (Fig. 2 J-L), without altering amplitude (Fig. S2 I, K), though overall this effect was not statistically significant. Of cells recorded, 54.5% showed an increase in GABA release in response to 5HT, 36.4% showed a decrease and 9.1% showed no change (Fig. S2L).

### Fear learning induces plasticity at vPAG^VGAT^-CeA inputs

We previously showed that chemogenetic inhibition of vPAG^VGAT^ neurons during fear learning impairs later fear expression [22], suggesting that the activity of these neurons during fear learning shapes conditioned fear processes. Because 5HT is released in the CeA during fear learning and modulates GABA release from vPAG^VGAT-^CeA inputs via 5HT_2C_ receptors, we hypothesized that fear learning may alter 5HT-induced GABA release from vPAG^VGAT^-CeA inputs via this mechanism. To test this hypothesis, we put intra-vPAG ChR2-expressing *Vgat*-ires-Cre mice through a single session of fear learning. Approximately 15 min after the session, we prepared slices containing the CeA for electrophysiology (Fig. 4 A-B). Naïve mice were treated identically but were presented with tones only. We then recorded the effects of 5HT (10 µM) on optically evoked eIPSCs from mice that underwent fear learning (fear) or control mice (naïve). Results showed that the ability of 5HT to enhance input-specific GABA release was blunted in fear mice relative to naïve mice (Fig. 4 C), evidenced by a significant change in the amplitude of the first evoked peak [F(14, 168)= 3.258, p= 0.0001 for the time x treatment interaction; F(14, 168)= 19.57, p < 0.0001 for main effect of time; F(1, 12)= 10.30, p= 0.0075 for main effect of treatment, two way RM ANOVA]. Post-hoc analyses revealed between-group differences at minutes 8, 10, and 13-14 [p<.05 for all Bonferroni multiple comparisons tests]. (Fig. 4C). Significant between-group differences in the effects of 5HT on the amplitude of the first eIPSC were observed [see Fig. 4D; F(1, 12)= 8.805, p= 0.0118 for fear x 5HT interaction; F(1, 12)= 30.94, p= 0.0001 for main effect of 5HT; F(1, 12)= 8.157, p= 0.0145 for main effect of fear, two-way RM ANOVA]. Post-hoc analyses show that 5HT significantly enhanced the amplitude of the first eIPSC relative to baseline values in naïve but not fear mice [p<.0001 Bonferroni post-hoc test for multiple comparisons, indicating that the ability of 5HT to enhance the amplitude of the first eIPSC was significantly blunted in fear mice. Taken together, these data may reflect an engagement of the vPAG^VGAT^-CeA pathway during fear learning, thereby occluding further release by serotonin in our bath application experiments.

**Figure 4.**
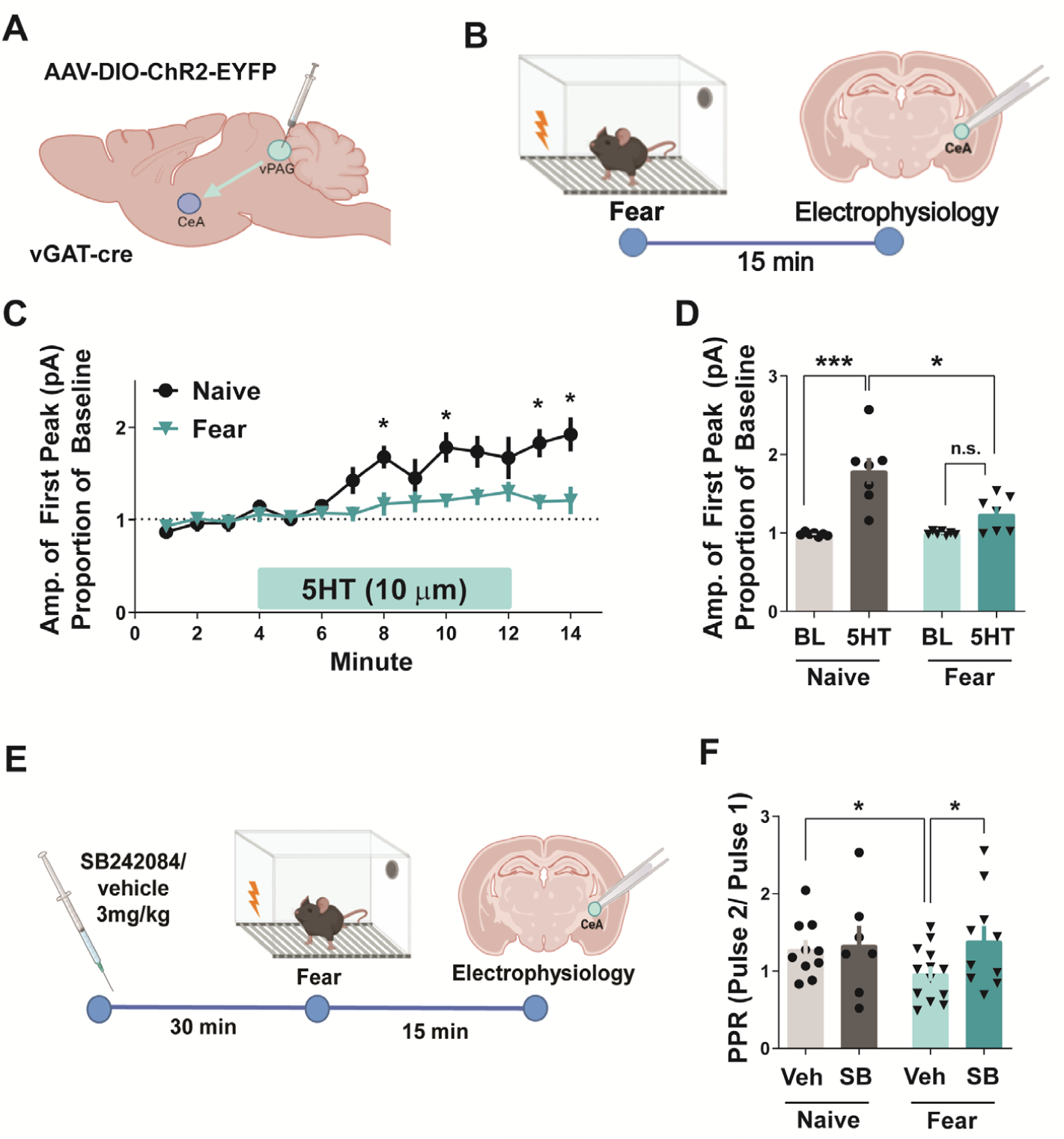
Fear learning engages the vPAG^Vgat^-CeA pathway. **A:** surgical schematic of channelrhodopsin infusion into the vPAG of *Vgat*-ires-Cre mice. **B:** Experimental timeline for fear conditioning and electrophysiology experiments. **C, D:** Time course and summary of the effect of bath application of serotonin (5HT; 10 µM) on the amplitude of the first evoked peak relative to baseline values in shock mice versus naïve mice (n= 7 cells per group from 3 mice per group), recordings in CeA. **E:** Experimental timeline for fear conditioning and electrophysiology experiments with 5HT_2c_ antagonist pretreatment. Mice were administered the 5HT2C antagonist SB242084 (3 mg/kg, i.p.) or vehicle prior to the fear acquisition session (shock) or 45 min prior to sacrifice (naïve) and recordings were performed at terminals in the CeA. **F:** Summary of the effects of serotonin on the paired pulse ratio (PPR; amplitude of pulse 2/ amplitude of pulse 1) in shock or naive mice that did or did not receive drug (**e**; n= 7-13 cells per group from 3-5 mice per group). *denotes *p*< 0.05; all data shown as means +/- SEM.

To examine this possibility, we compared PPR from mice that underwent fear learning (fear) or control mice (naïve) that were pretreated with 5HT_2C_ antagonist SB242084 or vehicle (Fig. 4E). We hypothesized that *in vivo* pretreatment with SB242084 would block engagement of 5HT_2C_ receptor signaling at the vPAG^VGAT^-CeA input during fear learning, focusing on the PPR as a measure that can be compared across *in vivo* conditions more readily than peak amplitude. Among mice that were not pretreated with SB24284, we found that fear mice had significantly lower PPRs than naïve mice [p= 0.0452, t(21)= 2.129 unpaired t-test] (Fig. 4F). Notably this effect of fear was blocked by pretreatment with SB24284, as fear mice pretreated with the drug had significantly higher PPRs than fear mice that received vehicle injections [p= 0.0476, t(21)= 2.104 unpaired t-test]. Pretreatment with SB242084 did not alter PPR in naïve mice relative to vehicle [p= 0.8286, t(15)= 0.2203 unpaired t-test]. These *a priori* hypotheses reflect between group differences, however a two-way ANOVA performed on this data set was not significant [F(1, 37)= 1.727, p= 0.1968 for the condition x drug interaction; F(1, 37)= 1.933, p= 0.1727 for the main effect of drug; F(1, 37)= 0.3345 for the main effect of condition].

### vPAG^VGAT^-CeA is dynamically engaged during fear and responds to shock-predicting cues

To characterize vPAG^VGAT^-CeA activity *in vivo*, we used fiber photometry coupled to the genetically encoded calcium indicator GCaMP7f to simultaneously record vPAG^VGAT^ cell bodies and terminals in the CeA during fear learning (Fig. 5A-C). We found that overall, cell body (vPAG) and terminal (CeA) activity was highly similar (Fig. 5 D, H). In mice that underwent fear conditioning, both vPAG and CeA responded to tone onset and shock, indicated by significant increases in average z-score compared to baseline [t(7)=3.843 p=.0063 for CeA tone response v BL, t(10)=4.473 p=.0012 for vPAG tone response v BL, t(7)=5.284, p=.0011 for CeA shock response v. BL, t(10)=11.31, p<.0001 for vPAG shock response v. BL] (Fig. 5D-G). Naïve mice did not show significant engagement of either region in response to tone onset (Fig. 5H-J), suggesting that vPAG^VGAT^-CeA responds specifically to cues that predict aversive stimuli, and not just neutral auditory cues.

**Figure 5.**
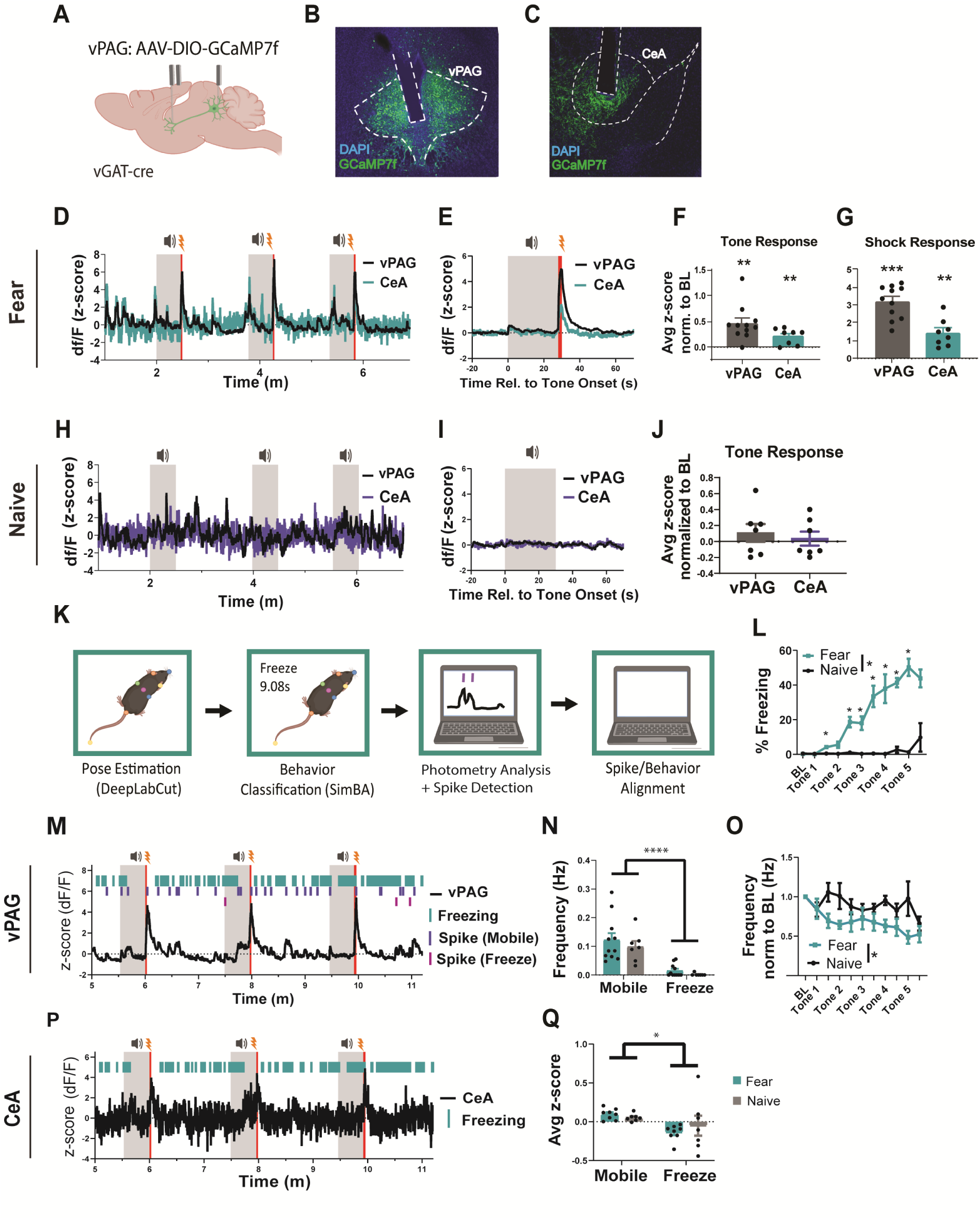
vPAG^Vgat^-CeA pathway is dynamically engaged during fear learning and responds to shock-predicting cues. **A:** surgical schematic of GCaMP7f infusion into the vPAG of *Vgat*-ires-Cre mice. **B, C:** representative image of GCaMP expression and fiber placement in vPAG (B) and CeA (C). **D:** Representative trace showing vPAG and CeA signal during fear learning in a mouse that underwent fear conditioning. **E:** vPAG and CeA responses to tone/shock presentation in fear conditioned mice averaged across cohort. **F:** Tone response calculated as the average from t=0-5, normalized to t= −5−0 relative to tone onset. **G:** Shock response calculated as the average from t=28-30, normalized to t=23−28 relative to tone onset. **H:** Representative trace showing vPAG and CeA signal during learning in a naïve mouse. **I:** vPAG and CeA responses to tone presentation in naïve mice averaged across cohort. **J:** Tone response calculated as the average from t=0-5, normalized to t= −5−0 relative to tone onset. **K:** schematic of analysis pipeline for alignment of freezing and photometry signal. **L**: Freezing behavior during fear learning. **M:** Representative trace showing alignment of freezing and vPAG signal with identified spikes classified as occurring during freezing (magenta) or mobility (purple). **N:** vPAG spike frequency during freezing and mobility for fear mice (turquoise) and naïve (gray). **O:** vPAG spike frequency during each epoch of fear learning, normalized to BL. **P:** Representative trace showing alignment of freezing and CeA signal. Q: average signal during freezing and mobility averaged across all bouts. n= 10 vPAG fear, 7 vPAG naïve, 8 CeA fear, 7 CeA naïve

Because our electrophysiology experiments indicate plasticity in the vPAG^VGAT^-CeA pathway after fear learning, we next asked whether differences in fear and naïve mice could be detected during fear expression. One day after fear learning, we returned mice to the same fear chamber and allowed them to forage for 15 minutes to characterize activity during context recall. Baseline activity was recorded for 5 minutes in a neutral, familiar environment prior to placing the mice back in the fear chamber in order to capture any shifts in activity. We did not find significant differences in responses between fear and naïve groups in either vPAG or CeA activity (Fig. S3 C-E). One day after context recall, mice underwent a cued recall test. Responses to tone presentations were quantified, and no differences were found in vPAG or CeA activity between groups (Fig S3 J-L).

### vPAG^VGAT^ neuron activity negatively correlates with freezing behavior

We previously showed that vPAG^VGAT^ neurons drive fear conditioning [22], and others have reported that vPAG^VGAT^ cells bidirectionally modulate freezing behavior [23]. However, it is unknown whether information about freezing is communicated via projections to the CeA. Therefore, we explored the relationship between vPAG^VGAT^-CeA activity and freezing by using machine learning-based algorithms to extract frame-by-frame behavior information (Fig. 5K). Comparison of photometry and behavior data revealed an inverse relationship between freezing and both vPAG^VGAT^ cell body and terminal activity.

During fear learning, fear mice froze significantly more than naïve mice, and had significantly lower vPAG spike frequency [freezing: two way RM ANOVA F(1,16)=56.82 p<.0001 main effect of fear, Spike frequency: F(1,16)=5.505 p=.0322 main effect of fear] (Fig. 5L, O). We next aligned freezing and vPAG spiking data to classify spikes as either occurring during freezing or mobility. We found that overall, vPAG spike frequency was significantly lower during freezing than mobility, indicating that vPAG activity is suppressed during freezing [two way ANOVA F(1,16)=52.49, p=.0001 for main effect of freezing, F(1,16)=.1275 p=.7257 for group x freezing interaction] (Fig. 5N). To determine whether vPAG^VGAT^ terminal activity is also suppressed during freezing, we extracted the start and end times of each freezing bout and parsed CeA data points as occurring during either freezing or mobility. We then took the average of each group. We chose to look at average activity in terminals because this method of calcium imaging is not sensitive enough to record spikes in terminals. Consistent with what we observed in cell bodies, activity in terminals is lower during freezing compared to mobility [two way ANOVA F(1,13)=7.959, p=.0144 for main effect of freezing] (Fig. 5Q). Notably, this effect was independent of whether or not mice underwent fear conditioning.

During context recall, fear mice exhibited higher freezing than naïve mice, but we did not find a difference in spike frequency across groups [freezing: F(1,16)= p=.0002 main effect of fear] (Fig S3). However, spike frequency in vPAG and average signal in CeA were significantly lower during freezing bouts, consistent with fear learning [vPAG: two-way ANOVA F(1,16)=69.84 p=.0001 main effect of freezing, CeA: two-way ANOVA F(1,12)=15.41 p=.002 main effect of freezing] (Fig S3 H,I). Similar effects were also present during cued recall. Fear mice froze more than naïve mice, had lower vPAG spike frequency, and lower activity during freezing compared to mobility [freezing: two way RM ANOVA F(1,16)=6.837 p=.0188 for main effect of fear, vPAG spike frequency: two way RM ANOVA F(1,16)=5.457 p=.0328 main effect of fear, vPAG spiking during freezing: F(1,15)=24.65 p=.0002 main effect of freezing, CeA : F(1,13)=23.38 p=.0003 main effect of freezing] (Fig. S3 M-P).

## DISCUSSION

The current experiments identify a GABAergic input to the CeA from the vPAG that is modulated by serotonin via presynaptic 5HT_2C_ receptors. We find that plasticity occurs in this pathway following fear learning and is mediated by 5HT_2C_. Together, these data demonstrate that the vPAG has inhibitory influence over the CeA and adds to a growing body of evidence that PAG projections to ‘upstream’ structures inform conditioned fear processes.

### Non-canonical ascending outputs from the vPAG-CeA

While descending amygdalar inputs to the PAG and their role in fear behavior have been well-established, reciprocal ascending inputs from the PAG to the amygdala are of growing interest [10,40]. Retrograde tracing studies have shown that cells of the vPAG innervate the CeA [18,19] but the cellular phenotype of these cells had not been characterized. The current results extend these findings by identifying GABA neurons as one population of vPAG neurons that robustly innervate the CeA. Using *in vivo* and *ex vivo* approaches, we confirmed that activation of the vPAG-CeA input promotes GABA release. Together, these findings demonstrate that the vPAG has inhibitory influence over the CeA and is anatomically and functionally poised to regulate CeA microcircuits and perhaps overall amygdalar output.

### Serotonin Dynamics in the CeA

Serotonin neurons in the DRN that project to the amygdala are engaged during fear learning and extinction [28,41], but the dynamics of 5HT release in the CeA has not been characterized. Microdialysis studies have shown that 5HT levels in the BLA are elevated after fear recall [8] and in both the BLA and CeA in response to noxious stimuli [42,43], but this technique can only measure serotonin levels on the scale of minutes. Additionally, DRN 5HT neurons co-express a variety of neurotransmitters in addition to serotonin [28,41,44], thus activity of DRN 5HT neurons does not necessarily reflect downstream release of 5HT. Recent advances in biosensor imaging allow for visualization of 5HT release with millisecond temporal resolution and high specificity, providing a level of detail not attainable with other methods.

We show for the first time that 5HT release in the CeA is dynamic during fear learning, consisting of both increases and decreases in extracellular levels. Additionally, we show that 5HT is released specifically in response to shock but not tone, and that release changes over the course of fear learning. Previous studies show that CeA-projecting 5HT cell bodies in the DRN are activated by shock [28], consistent with the upward response we observe after shock end. However, the downward spike during shock has not been documented before and may be driven by modulation of DR 5HT terminals.

### Serotonin modulation of GABAergic signaling in CEA

Our results show that serotonin robustly and reliably enhances presynaptic GABA release from the vPAG-CeA input, observed as enhanced frequency, but not amplitude, of GABAergic events. These observations indicate that serotonin enhances presynaptic GABA transmission without effects on post-synaptic transmission. Additionally, this effect persisted in the presence of TTX, demonstrating that modulation of GABA release at this input is direct and does not require an intervening network. We also found evidence that serotonin enhances GABA release in the general medial CeA population, but unlike the input-specific effect, 5HT-mediated enhancement of GABA release was reduced in the presence of TTX. As such, serotonin’s modulation of GABA release within the CeA appears to be more varied in the general population. Together, these results reveal a novel and complex role for serotonin in modulating inhibitory transmission within the CeA.

Based on observations that the 5HT_2C_ receptor modulates GABA release in the CeA [32], we hypothesized that serotonin’s effects in the CeA may involve the 5HT_2C_ type receptor. In agreement with this hypothesis, we found that 5HT enhancement of GABA release from the vPAG-CeA input was abolished by a 5HT_2C_ antagonist. Because 5HT_2C_R blockade prevented this effect, measured via PPR, we conclude that these receptors have a pre-synaptic locus of action. Generally, 5HT_2C_ receptors are thought to be located at post-synaptic sites, where they control cellular excitability [39]. However, a pre-synaptic locus of action has been previously shown, evidenced by findings that receptors co-localize with pre-synaptic proteins [46] and modulate GABA release in the VTA in a calcium-dependent manner [31]. Thus, pre-synaptic 5HT_2C_ receptors that control neurotransmitter release do not appear to be unique to the CeA. In contrast, we found that 5HT enhancement of GABA transmission in the general CeA is only partially blocked by 5HT_2C_ receptors, suggesting that additional 5HT receptors modulate GABA release in the greater CeA. Therefore, serotonin may finely tune inhibitory transmission within the CeA in a synapse-specific manner, with 5HT_2C_ receptors restricted to select synaptic nodes like those arising from vPAG GABA inputs.

### A locus for 5HT modulation of fear learning in the CeA

Fear learning requires plasticity at specific nodes to encode associations between fear-inducing stimuli and the environmental cues that predict them. Within the CeA, adaptations in excitatory transmission are observed following fear learning that are likely driven by glutamatergic inputs from the BLA, lateral amygdala, paraventricular nucleus of the thalamus, or cortex [12–14,47]. Enhancements in inhibitory modulation have also been reported, though from unidentified sources [48]. The current findings suggest that plasticity at the vPAG-CeA input may be an additional adaptation underlying fear learning. Specifically, we observed that GABA release from this input was augmented following fear learning.

Additionally, serotonin’s ability to further facilitate GABA release from this input was disrupted following fear learning, suggesting that plasticity had occurred at the vPAG-CeA input. This finding is consistent with the hypothesis that the serotonin-GABA system is engaged endogenously during fear learning, thereby occluding the ability of 5HT to elicit further GABA release in slice recordings. We confirmed this possibility by blocking 5HT_2C_ receptors *in vivo* prior to fear learning, which resulted in normalized GABA release from the vPAG-CeA input. Based on these findings and the model revealed through characterization of the vPAG-CeA input, we posit that serotonin activates presynaptic 5HT_2C_ receptors in the medial CeA during fear learning, which stimulates GABA release from the vPAG-CeA input (Fig. S4). As such, it appears that serotonin recruits the vPAG’s inhibitory modulation of the medial CeA.

### Dynamic Engagement of the vPAG^VGAT^-CEA pathway in fear learning

It has been previously shown that vPAG GABA neurons bidirectionally modulate freezing behavior, and optogenetic inhibition and activation of GABA neurons increase and decrease freezing behavior respectively [23]. Consistent with this, we found that activity in vPAG^VGAT^ cell bodies and terminals in CeA is lower during freezing than mobility. However, this effect was independent of whether mice underwent fear learning or not and thus suggests that vPAG^VGAT^ neurons modulate freezing behavior under basal conditions, a property that is not altered by fear learning. Notably, our findings suggest an additional role for vPAG^VGAT^ cells in fear learning beyond modulation of freezing. We previously reported that chemogenetic inhibition of vPAG^VGAT^ cells during fear learning impaired subsequent fear expression [22], and in the current study, we show that fear learning induces plasticity in vPAG projections to CeA. Taken together, this suggests fear-specific alterations in signaling in this pathway. To test this hypothesis, we used fiber photometry to record VGAT cell bodies in vPAG and terminals in CeA during fear learning and recall. We found that vPAG^VGAT^ cell bodies and terminals in CeA were responsive to both tone and shock in fear mice. Importantly, cell bodies and terminals were not responsive to tone in naïve mice, suggesting that vPAG^VGAT^ cells are not merely responsive to auditory cues, but specifically to cues that predict harm. The notion that vPAG^VGAT^-CeA might play a role in the prediction of aversive stimuli is consistent with recent findings that ventrolateral PAG neurons encode the size of a threat [49,50]. Together, these findings implicate the vPAG in more complex fear encoding processes than previously thought. Additionally, we did not see differences in vPAG^VGAT^-CeA activity between fear and naïve mice during context or cued recall, indicating that this pathway is important for the acquisition but not expression of conditioned fear.

### Using automated tracking to identify relationships between behavior and cellular activity

Recent advances in open-source rodent behavior tracking technology allows for fast, precise, and accurate identification of specific behaviors. Machine learning-based platforms like DeepLabCut and Simba make it possible to extract frame-by frame behavior data from recorded videos, allowing for discrete neural events to be mapped onto corresponding behavior with high temporal precision. An advantage of using these platforms is that models can be easily developed to identify any behavior of interest, thus they are highly flexible and can be used for a variety of applications. Here, we develop an analysis pipeline that allows us to compare freezing to photometry data in two different ways; by aligning spiking, and by calculating average activity during different behavioral states. Using the spike alignment method, we show that vPAG^VGAT^ spike frequency is lower during freezing compared to mobility. Complementary to this, we find the same effect in vPAG terminals in the CeA using the averaging method. These techniques provide a framework for parsing fiber photometry recordings that can accommodate different types of *in vivo* imaging data.

In conclusion, our study characterizes a novel GABAergic input to the CeA from the vPAG that is engaged during fear learning and is modulated by presynaptic 5HT_2C_. These findings provide initial evidence that the vPAG has modulatory influence over fear learning through inputs to the CeA. As such, the vPAG is not a mere effector that executes commands from the CeA to drive fear expression but rather may shape the learning processes that underlie these responses. Given the reciprocal interactions between the vPAG and CeA and their established roles in fear learning and expression, dysregulation of the vPAG^VGAT^-CeA input may contribute to the pathophysiology of post-traumatic stress and other fear-related disorders and therefore may be a novel target for therapeutic interventions.

## ACKNOWLEDGEMENTS

Schematics for this manuscript were created using Biorender.com

## AUTHOR CONTRIBUTIONS

O.J.H, E.G.L.G. and T.L.K. designed the experiments. O.J.H, E.G.L.G., J.F.D., J.S., D.W.B., M.H., A.K., N.M.M. and A.J.L. performed the experiments. O.J.H, E.G.L.G., J.F.D., C.M.M. and J.A.H. analyzed data. O.J.H, E.G.L.G. and T.L.K. interpreted the data and wrote the manuscript.

## Supplementary Information

**Figure S1.**
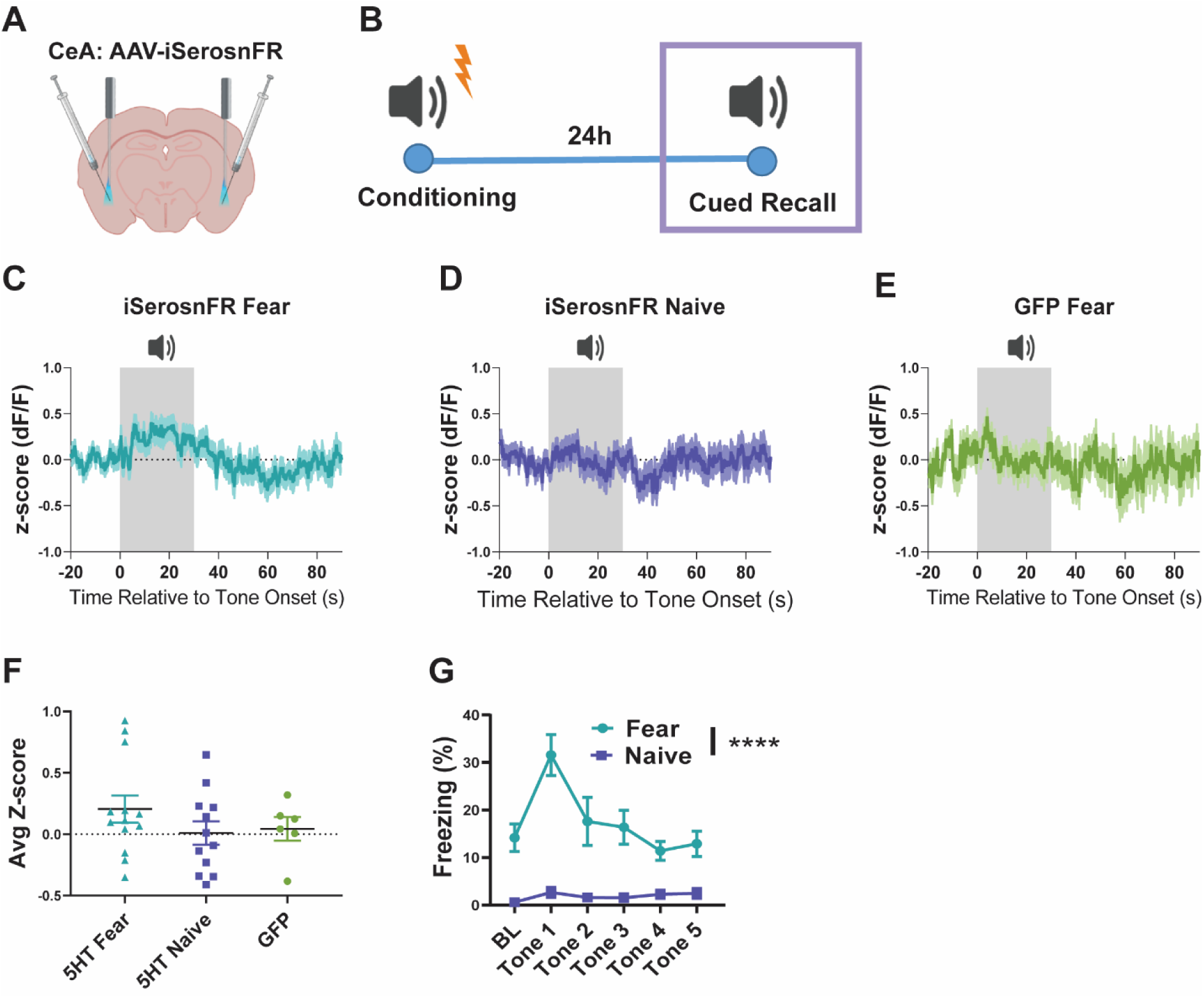
5HT release during cued fear recall. **A:** surgical schematic of iSeroSnFR and fiber implant in CeA. **B:** Experimental timeline for fear conditioning. **C-E**: responses to tone presentation during fear recall averaged across groups for iSeroSnFR fear (C), iSeroSnFR naïve (D) and GFP fear (E). **F:** tone response averaged from t=0-5s relative to tone onset across all trials. **G:** freezing behavior during cued recall. n= 13 iSeroSnFR fear, 11 iSeroSnFR naïve, 6 GFP fear

**Figure S2.**
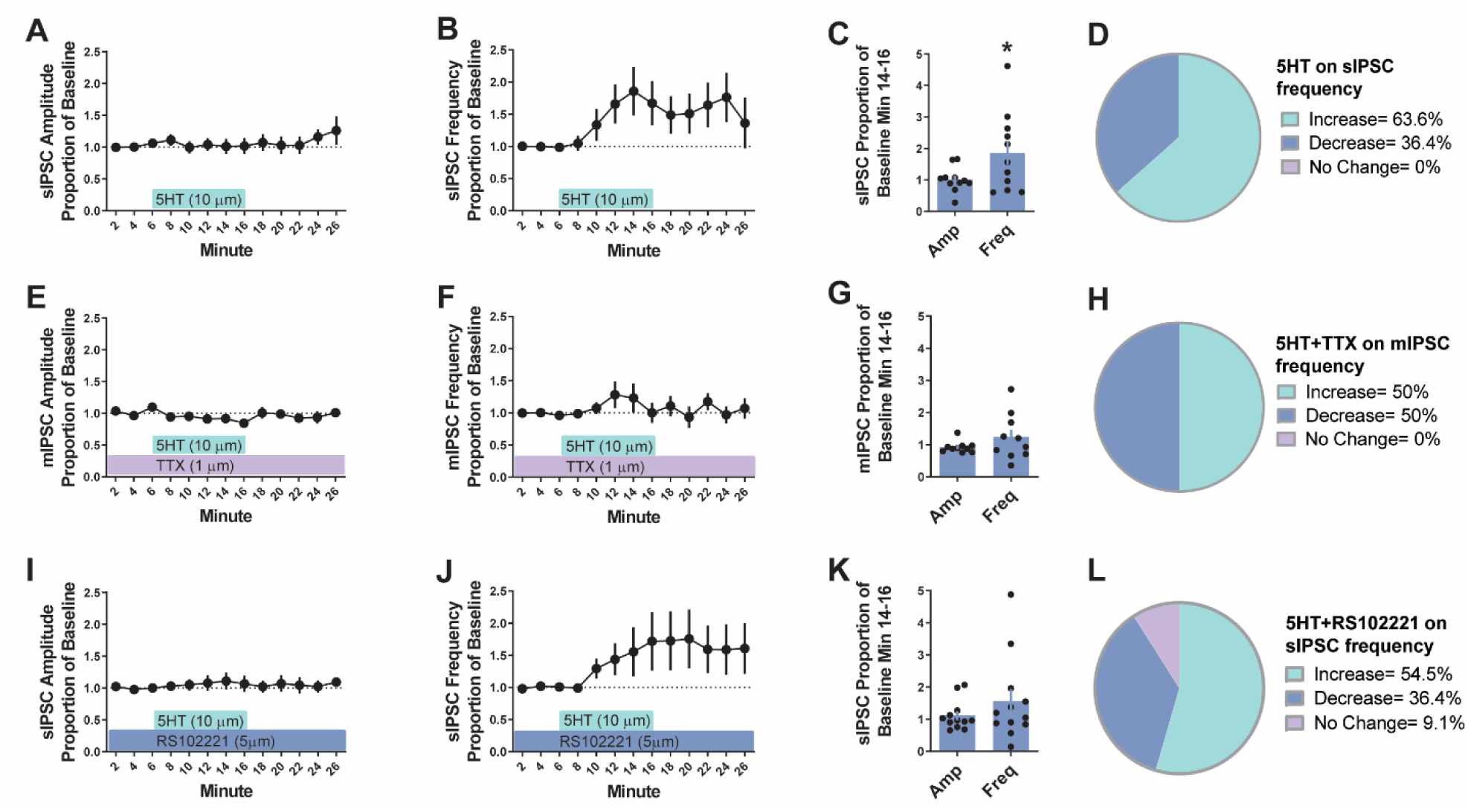
Serotonin facilitates pre-synaptic GABA transmission in the CeA. **A-B:** Time course of the effects of bath application of serotonin (5HT; 10 µM) on the amplitude (A) and frequency (B) of spontaneous inhibitory post-synaptic currents (sIPSC) relative to baseline values (n= 11 cells from 7 naïve mice). **C**: Summary of the effects of serotonin on sIPSC amplitude and frequency. **D:** proportion of cells showing different effects of 5HT on sIPSC frequency in the greater medial CeA. **E-F:** Time course of the effects of bath application of serotonin (10 µM) on the amplitude (E) and frequency (F) of miniature inhibitory post-synaptic currents (mIPSC) relative to baseline values in the presence of TTX (1 µM) (n= 10 cells from 7 naïve mice). **G:** Summary of the effects of serotonin on mIPSC amplitude and frequency in the presence of TTX. **H:** proportion of cells showing different effects of 5HT on sIPSC frequency in the presence of TTX (1 µM). **I-J**: Time course of the effects of bath application of serotonin (10 µM) on the amplitude (I) and frequency (J) of sIPSC relative to baseline values in the presence of the 5HT2C receptor antagonist RS102221 (5 µM) (n= 11 cells from 6 naïve mice). **K**: Summary of the effects of serotonin on sIPSC amplitude and frequency in the presence of RS102221. **L:** proportion of cells showing different effects of 5HT on sIPSC frequency in the presence of RS102221 (5 µM). * denotes *p*< 0.05; all data shown as means +/- SEM. Summaries were based on PPR data for vPAG-CeA input effects and IPSC frequency data for effects in the greater CeA population. Increased release was defined as >10% of baseline values; decreased release was defined as <10% of baseline values; no change was defined as within +/- 10% of baseline values.

**Figure S3.**
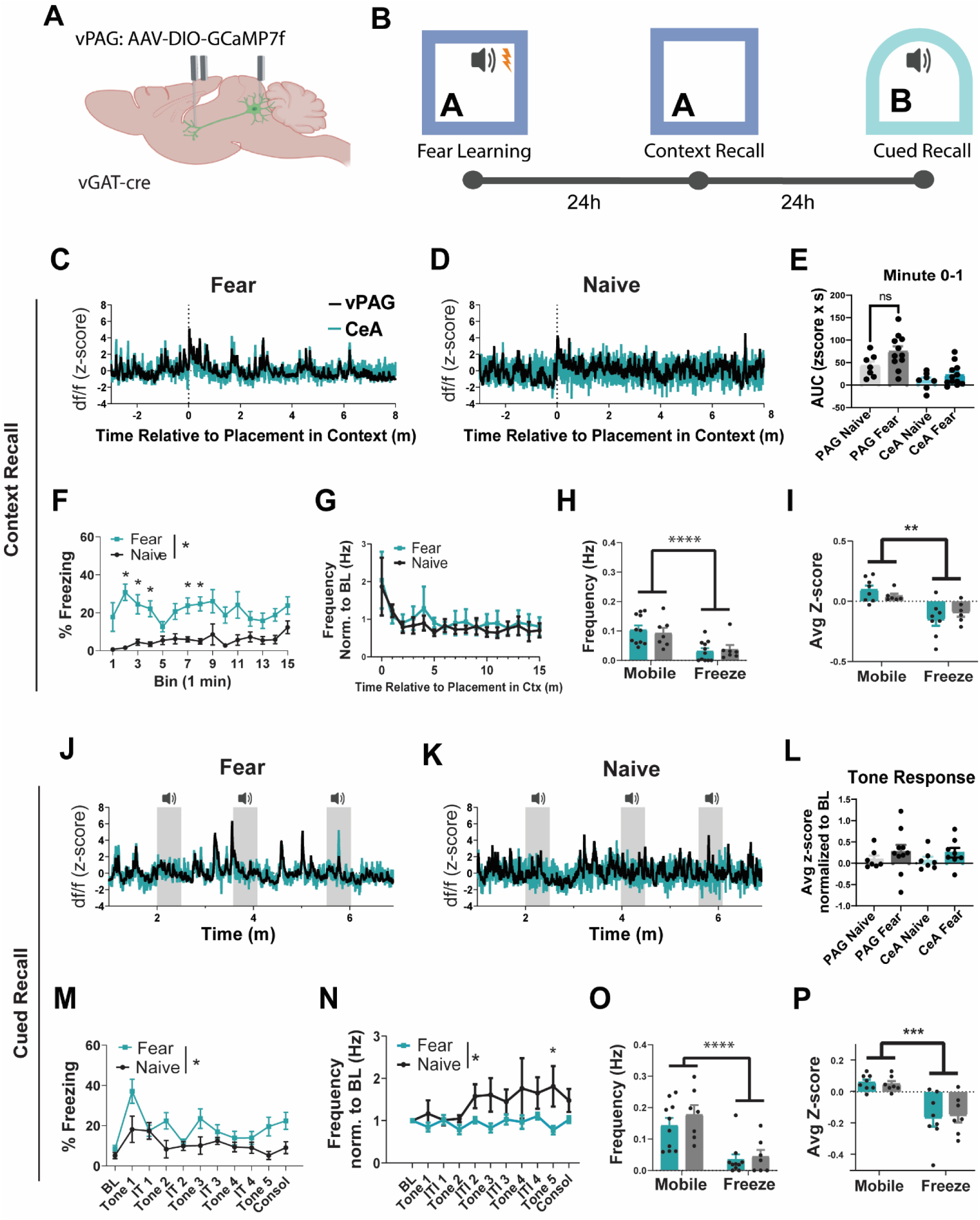
vPAG-CeA dynamics during fear recall. **A:** surgical schematic of GCaMP7f infusion into the vPAG of *Vgat*-ires-Cre mice. **B:** Experimental timeline for fear learning and recall. **C-D:** Representative traces from a fear conditioned (C) and naïve (D) mouse during context recall. **E:** Area under curve for the first minute after the mouse is placed into the fear context. **F:** Freezing behavior during context recall. **G:** vPAG spike frequency in 1 minute bins. **H**: vPAG spike frequency during freezing and mobility averaged across all bouts. **I:** CeA signal during freezing and mobility averaged across all bouts during context recall. **J-K:** Representative traces from a fear conditioned (J) and naïve (K) mouse during cued recall. **L:** Tone response calculated as the average from t=0-5 normalized to t=-5-0 relative to tone onset. **M:** Freezing behavior during cued recall. **N:** vPAG spike frequency during each epoch of cued recall. **O**: vPAG spike frequency during freezing and mobility averaged across all bouts. **P:** CeA signal during freezing and mobility averaged across all bouts during cued recall. n= 10 vPAG fear, 7 vPAG naïve, 8 CeA fear, 7 CeA naïve.

**Figure S4.**
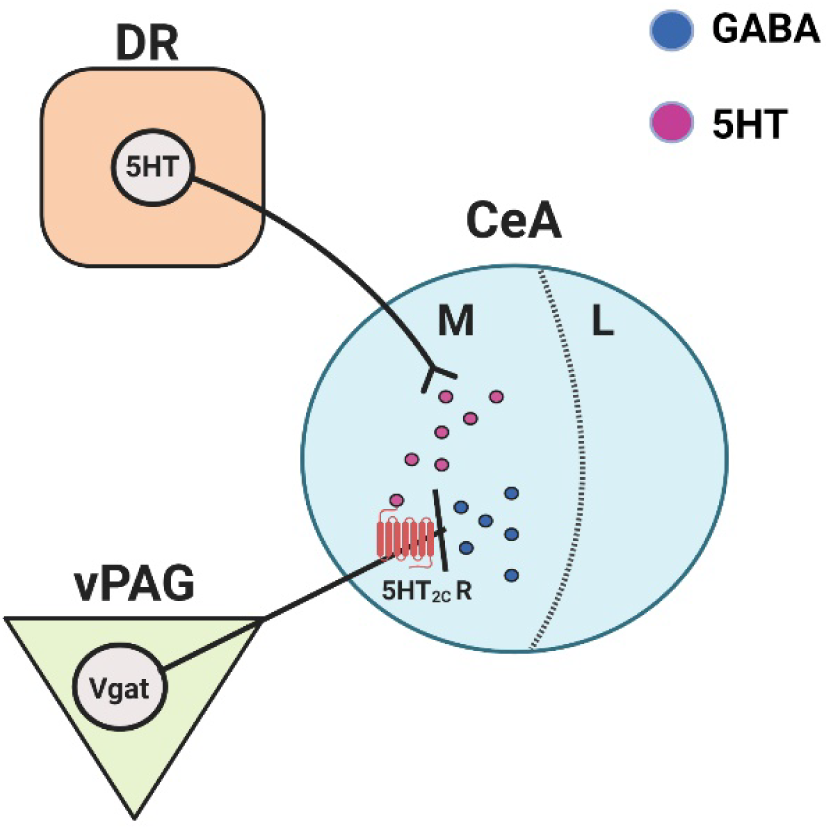
Working model of 5HT modulation of GABA transmission via presynaptic 5HT_2C_ in the vPAG^VGAT^-CeA pathway. vPAG GABA neurons project to the CeA. 5HT released from dorsal raphe inputs acts on 5HT_2C_ receptors located on vPAG^VGAT^ terminals to potentiate GABA release in the CeA. Fear learning induces plasticity in the vPAG^VGAT^-CeA pathway in a 5HT_2C_ receptor dependent manner.

